# Mutations on RBD of SARS-CoV-2 Omicron variant result in stronger binding to human ACE2 receptor

**DOI:** 10.1101/2021.12.10.472102

**Authors:** Cecylia S. Lupala, Yongjin Ye, Hong Chen, Xiao-Dong Su, Haiguang Liu

**Author notes:** Corresponding authors: Xiao-Dong Su, Or Haiguang Liu.

## Abstract

The COVID-19 pandemic caused by the SARS-CoV-2 virus has led to more than 270 million infections and 5.3 million of deaths worldwide. Several major variants of SARS-CoV-2 have emerged and posed challenges in controlling the pandemic. The recently occurred Omicron variant raised serious concerns about reducing the efficacy of vaccines and neutralization antibodies due to its vast mutations. We have modelled the complex structure of the human ACE2 protein and the receptor binding domain (RBD) of Omicron Spike protein (S-protein), and conducted atomistic molecular dynamics simulations to study the binding interactions. The analysis shows that the Omicron RBD binds more strongly to the human ACE2 protein than the original strain. The mutations at the ACE2-RBD interface enhance the tight binding by increasing hydrogen bonding interaction and enlarging buried solvent accessible surface area.

## Introduction

The COVID-19 pandemic caused by the SARS-CoV-2 is affecting global health and economy seriously[1]. According to JHU CSSE COVID-19 Data[2], there are 270 million infections and over 5.3 million fatalities as of December 13, 2021. Several vaccines have been developed and applied to prevent the spreading of SARS-CoV-2 viruses[3], however, these efforts are challenged by emerged virus variants due to mutations [4–7]. Among major variants, several strains were called out to be ‘variant of concerns (VOC)’ by the world health organization (WHO). On November 26, 2021, the WHO named a new variant (B.1.1.529) to be Omicron, designated to be a VOC [8]. The Omicron variant has accumulated a vast number of mutations, particularly in spike protein that is responsible for the initiation of infection through cell entry. There are 15 mutations on the receptor binding domain (RBD) of the spike protein, which has over 30 mutations in total (see Figure 1) [8,9]. Such a large number of accumulated mutations is unprecedent. Because the spike protein is not only the receptor ACE2 (Angiotensin converting enzyme 2) binding partner [10,11], but also the major antigenicity site, thus the target of many antibodies or drugs, it is crucial to investigate the impacts to the efficacy of neutralizing antibodies, under the concerns of immune escapes. Furthermore, about 10 mutations occur at the RBD binding interface to the ACE2 receptor protein. This level of mutation also raised a serious question on how the RBD of Omicron variant binds to the ACE2. Will the binding become stronger or weaker, and whether there is a need for an alternative receptor to facilitate the infection of human cells?

**Figure 1.**
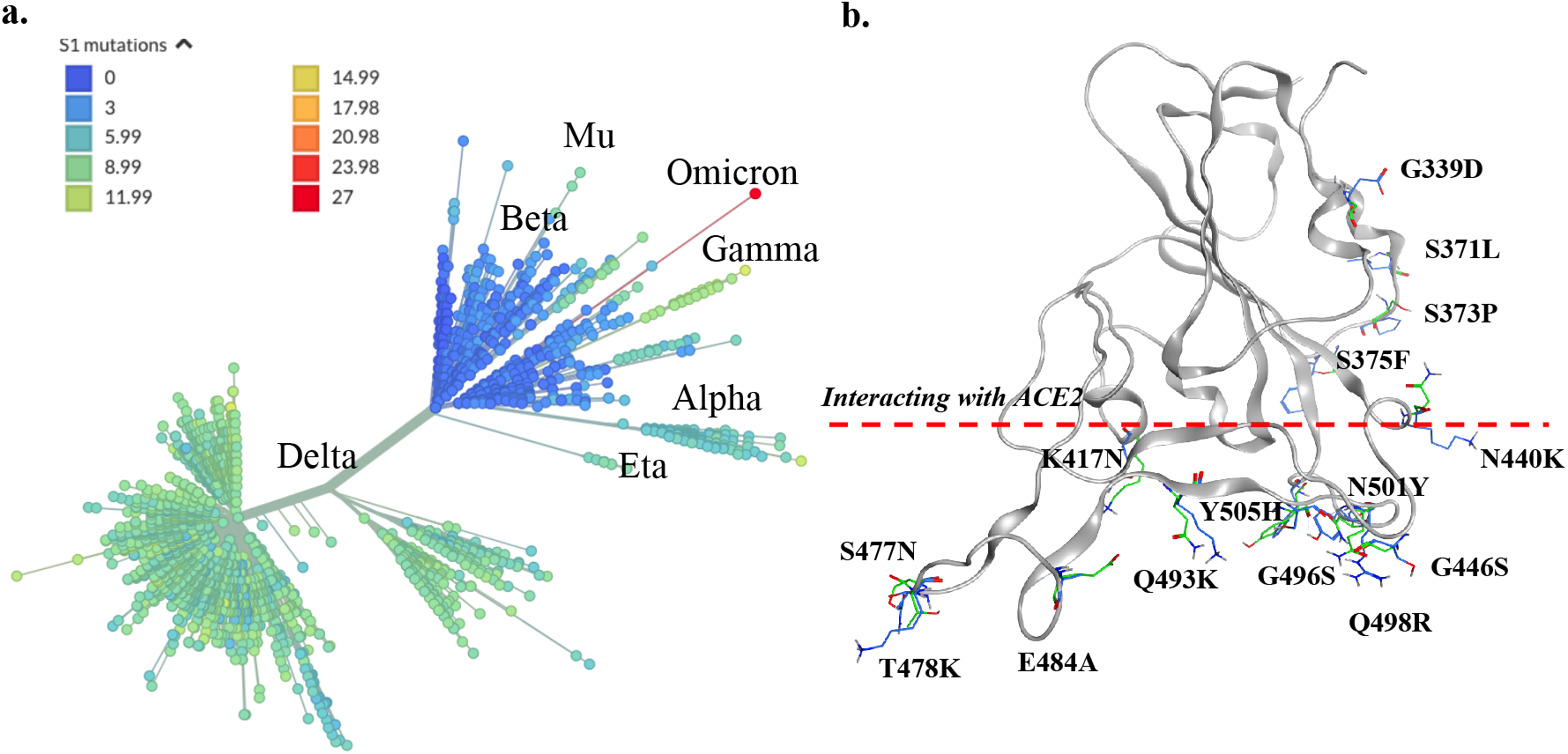
Mutations and the diversity of SARS-CoV-2. (**a**) The phylogenetic tree of SARS-CoV-2. Major variants are labelled on the graph, and the color of clans is according to the number of spike protein mutations. The tree is generated at https://nextstrain.org. (**b**) Mutation sites of the receptor binding domain. The residues below the red line are at or near the ACE2 binding interface.

Computational modeling and dynamics simulations have been applied to investigate the interactions between the SARS-CoV-2 RBD and the ACE2 receptor [12,13]. Before the structures of RBD-ACE2 complex were resolved experimentally, homology modeling and simulations have successfully predicted the model and quantified the interactions [13,14]. Computer simulations were also used to study the interactions between RBD and ACE2 from other mammals, and the results provide hints on molecular mechanism for SARS-CoV-2 infection to other animals [15,16]. Here, we followed a similar approach, constructed the structure of human ACE2 and the RBD of Omicron variant (hereafter denoted as ACE2-RBD^O^, where the superscript indicates Omicron). Then the complex structure was subjected to atomistic molecular dynamics simulations to refine the model and to probe the dynamical interactions between ACE2 and RBD. After comparing to the wild type ACE2-RBD complex system, we found that the RBD^O^ exhibits stronger binding to human ACE2, suggesting that the Omicron variant infects cells via the same mechanism and the infectivity might be enhanced due to the stronger binding interactions.

## Results

### The structures of Omicron RBD and ACE2-RBD complex are stable

The averaged backbone root-mean-square-deviation (RMSD) of the RBD is less than 1.4 Å compared to the starting model for both the wild type and Omicron systems (Figure 2a). For the wild type RBD, the structure ensembles from two independent simulations (each 500 ns) deviated from the crystal structure of RBD by 1.2 Å on average; interestingly, the RBD^O^ has averaged RMSD values of 1.4 Å, indicating that the mutations only slightly alter the structure of RBD^O^ from the wild type RBD. Similarly, the ACE2-RBD complexes are stable through simulations, reflecting on the RMSD with respect to starting complex structures (Figure 2b). The RMSD for the wild type ACE2-RBD complex is averaged to 3.0 Å and 2.5 Å for the structures sampled from the two trajectories; while the values are 2.2 Å and 2.6 Å for the two simulation trajectories of ACE2-RBD^O^. Therefore, we predict that the mutations in Omicron variant do not significantly reduce the RBD stability, instead, the ACE2-RBD^O^ complex is even slightly more stable than the wild type, according to the RMSD analysis. The residue fluctuations were analyzed by calculating the root-mean-square-fluctuations (RMSF) of the RBD (Figure 2c). According to the average values of RMSF, the RBD^O^ is more rigid than its wild type (1.5 Å vs. 2.1 Å). The reduction of the RMSF is more pronounced at the interfacing residues of RBD, also known as the receptor binding motif (RBD, residues 434-508) [17]. We also closely examined the fluctuations of mutated residues (Figure 2c, right panel) and found that the 15 mutated residues in the Omicron variant consistently exhibit smaller fluctuations, compared to their wild type counterparts. It is plausible that the binding of ACE2 stabilize these residues, which in turn enhance the stability of the ACE2-RBD^O^ complex. Detailed quantifications on interactions between ACE2 and RBD are elaborated in the following sections.

**Figure 2.**
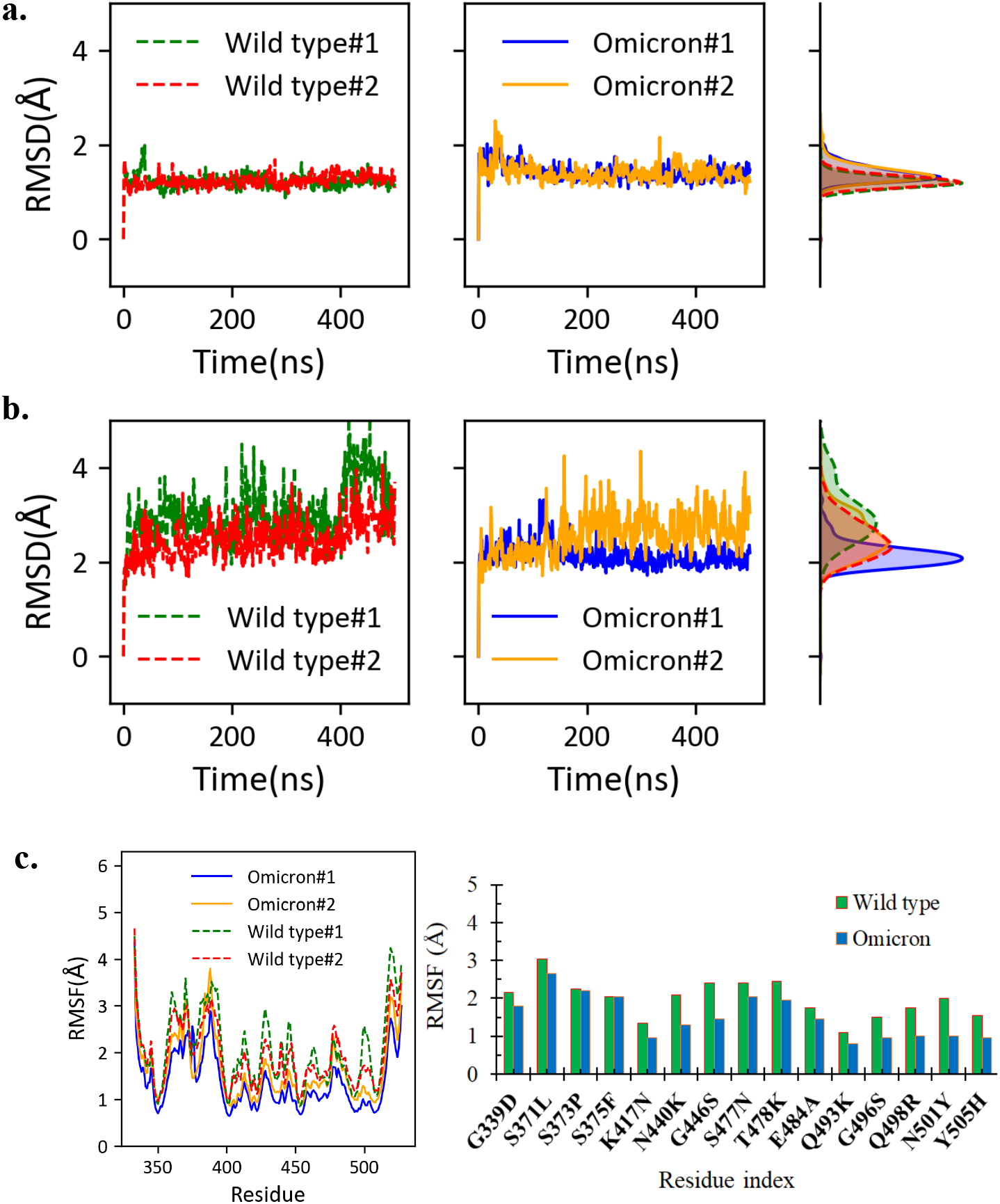
Stability of the RBD and ACE2-RBD complex structures. (**a**) The RMSD of RBD with respect to the starting structure. The histogram of each RMSD time trace is drawn on the right. (**b**) The RMSD of the whole complex with respect to the starting complex structure, with the histograms shown on the right. (**c**) The RBD residue fluctuations. The residue fluctuations for the mutation sites are shown on the right panel.

### The interactions between ACE2 and RBD are enhanced in Omicron variant

We extracted the hydrogen bonds formed directly between ACE2 and RBD^O^, and further compared the data to the wild type system (Figure 3). On average, there are 6.5 ± 2.2 hydrogen bonds formed between ACE2 and RBD^O^, about 10% more than 5.9 ± 2.4 hydrogen bonds observed in the wild type system. A closer examination on the specific hydrogen bonds reveals that the Q493K and N501Y play important roles in forming new hydrogen bonds (Table 1). It is worthwhile to note that the hydrogen bonds are very dynamical, and the total number of hydrogen bonds at any instant time fluctuates significantly. Therefore, in the table we only listed seven hydrogen bonds that are frequently observed during simulations, with the occupancy close to 20% or above. As shown in Table 1, the only hydrogen bond with occupancy below 20% is between ACE2 S19 and RBD A475 (occupancy = 18.73%). In the case of ACE2-RBD^O^, the next frequently observed hydrogen bond is between K31 of ACE2 and W456 of RBD^O^ with an occupancy of 16.25% (not listed in Table 1). As shown in Table 1, there are five common stable hydrogen bonds observed in both the wild type and Omicron variant systems. The mutations resulted in the loss of two hydrogen bonds: (1) the K417N mutation caused the loss of hydrogen bonding with ACE2 residue D30, and (2) the Y505H mutation significantly reduced its bonding to E37 of ACE2. The Q493K mutation not only maintains the hydrogen bond between Q493 and E35 of ACE2 in the wild type complex, but also adds the possibility of forming a new stable hydrogen bond between K493 and the D38 of ACE2. The hydrogen bond between Y501 of RBD^O^ and the Y41 of the ACE2 is also a new hydrogen bond frequently observed in simulations. The hydrogen bond between the S19 of ACE2 and the A475 of RBD^O^ is stronger than that in the wild type system, although neither residues were mutated in the Omicron variant. It is possibly influenced by the local changes due to the S477N and T478K mutations. By comparing the occupancies, we conclude that the hydrogen bonds between ACE2 and RBD^O^ are more stable through the simulations, and therefore resulting more hydrogen bonds on average.

**Figure 3.**
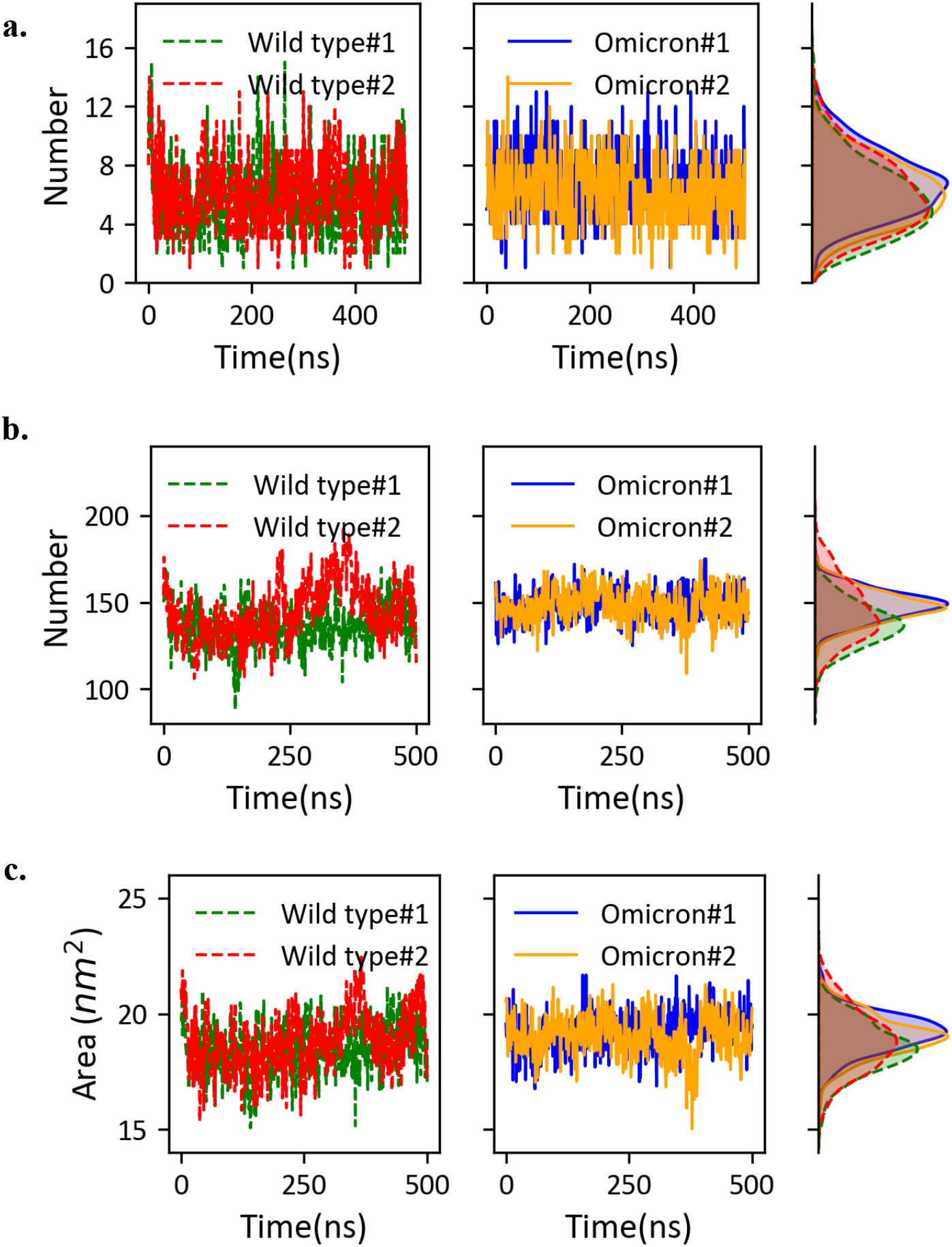
Quantitative analysis of the ACE2-RBD interactions. (**a**) Hydrogen bonds between ACE2 and RBD/RBD^O^. The time traces of hydrogen bond numbers observed during the simulations are shown on the left and middle columns. The histograms are shown on the right column to compare the statistics between the wild type system and the Omicron variant system. (**b**) The number of residue contacts between ACE2 and RBD. (**c**) The buried surface area due to ACE2-RBD binding. Similar to (a), the histograms are shown to facilitate the comparison in (b, c).

**Table 1.** Hydrogen bonds between the RBD and the ACE2.

| Wild type ACE2-RBD |  |  | Omicron ACE2-RBD |  |  |
| --- | --- | --- | --- | --- | --- |
| ACE2 | RBD | Occupancy | ACE2 | RBD<sup>o</sup> | Occupancy |
| K353-Main | G502-Main | 57.97% | E35-Side | K493-Side<br>* | 75.67% |
| Y83-Side | N487-Side | 50.60% | D355-Side | T500-Side | 60.52% |
| E35-Side | Q493-Side | 38.84% | K353-Main | G502-Main | 57.03% |
| D30-Side | K417-Side | 34.36% | D38-Side | K493-Side<br>* | 52.34% |
| D355-Side | T500-Side | 25.60% | Y83-Side | N487-Side | 52.04% |
| E37-Side | Y505-Side | 21.71% | S19-Side | A475-Main | 36.59% |
| S19-Side | A475-Main | 18.73% | Y41-Side | Y501-Side<br>* | 35.00% |

\* The residues were mutated from the wild type RBD
The entries shaded in blue color are either not lost or with low occupancy in the Omicron variant system; the entries with yellow shading are the new hydrogen bonds observed in the Omicron variant; the other entries are the common hydrogen bonds in both wild type and Omicron systems.

We computed the number of van der Waals contacts between the ACE2 and RBD, as well as the buried surface area, to further assess the interactions between ACE2 and RBD. For the wild type system, the two simulations yield 137 ± 12 contacts on average, while the ACE2-RBD^O^ has 148 ± 9 contacts on average (Figure 3b). The statistics on the buried surface areas are consistent with the level of contacts. The Omicron variant resulted an increase of buried surface area from 18.5 nm^2^ to 19.1 nm^2^ (Figure 3c).

### The representative structures are highly similar

The representative structures were selected from the most populated clusters for the ACE2-RBD complexes. The largest cluster accounts for about 19.8% of the simulated structures for the wild type complex, and the largest cluster for the Omicron complex accounts for 38.4% of the sampled structures. The RBD structures are similar in the representative models, both within 1.4 Å backbone RMSD from the crystal structure (see Figure 4). In particular, the RBM regions are aligned very nicely (with backbone RMSD < 0.5 Å) for these structures, in accordance with the tight binding to ACE2. We computed the electrostatic potentials by solving the Poisson-Boltzmann equation for the RBM region in three structures (Figure 4b): the crystal structure and representative structure of the wild type RBD, as well as the representative structure of the RBD^O^. For the wild type RBD, positive and negative potential patches are dispersedly located at the binding interface. Strikingly, the same interface has larger patches with positive potentials in the RDB^O^. For instance, the region around G446S-Q493K-G496S-Q498R-N501Y-Y505H mutation sites exhibits stronger positive electrostatic potentials, improving its complementary to the charge surface of ACE2 protein (Figure 4c). In the corresponding region, the key residues from ACE2 are composed of D38-Y41-Q42-D355-S446, forming a negatively charged patch. We computed the binding energies for the representative models. In this case, we obtained one representative structure from each simulation trajectory using the same clustering algorithm, then we obtained two representative structures for the wild type ACE-RBD, and two for the Omicron variant system. The binding energies for the two wild type ACE2-RBD structures are −104.17 kcal/mol and −97.73 kcal/mol. The binding energies for ACE2-RBD^O^ structures are even lower (−112.25 kcal/mol and −107.04 kcal/mol), indicating stronger binding between ACE2 and RBD^O^.

**Figure 4.**
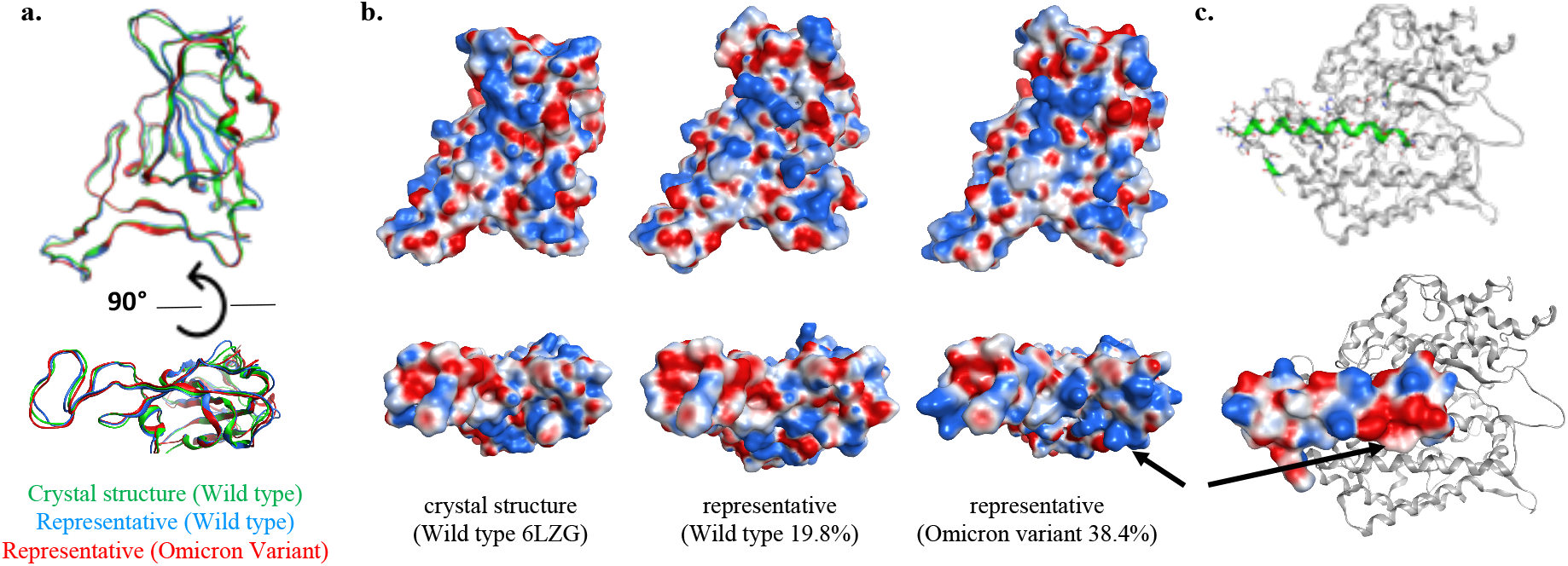
Representative structures and the electrostatic potential surfaces. (**a**) The representative structures of wild type RBD (blue) and RBD^O^ (red) are superposed to the crystal structure (green). The bottom panel shows the structure alignment for the ACE2 binding interface of RBD. (**b**) The electrostatic potentials on the RBD/RBD^O^ surface (−5 k_b_T/e to +5 k_b_T/e, for colors from red to blue). (**c**) The RBD binding interface of ACE2 and its electrostatic potentials (calculated from the crystal structure of ACE2). The black arrows point to the largest positive (on RBD^O^) and negative potential patches (on ACE2).

### Detailed structure features at the ACE2-RBD interface

The interactions at the interface of ACE2-RBD complex for the wild type have been previous reported in the perspectives of both static crystal structures [18,19] and dynamical conformations [13]. Generally, ACE2 residues 19-42 of the N-terminal helix, 82-83 near the η1, N330 at helix-13 and 352-357 at the β-hairpin-4,5 are in close contacts with RBD. For the RBD, crystal structures show that residues K417, G446, Y449, Y453, L455, F456, A475, F486, N487, Y489, Q493, Y495, G496, Q498, T500, N501, G502 and Y505 form direct contacts with human ACE2, while simulations have revealed additional residues Q474, G476, S477, T478, E484 and G485 at the loop (L_67_) of RBD to enhance the interactions [13]. Out of the 15 RBD mutations found in the Omicron variant, 10 residues (K417N, N440K, G446S, S477N, T478K, E484A, Q493K, G496S, Q498R, N501Y and Y505H) are located at ACE2-RBD interface, consequently changing the electrostatics surface charges at the interface and may have additional effects on the binding of antibodies and drugs targeting the interface due to the bulkier size of the mutant sidechains such as in T478K. This also applies for the mutant residue N440K at a loop near the binding interface with ACE2 (see Figure 5).

**Figure 5.**
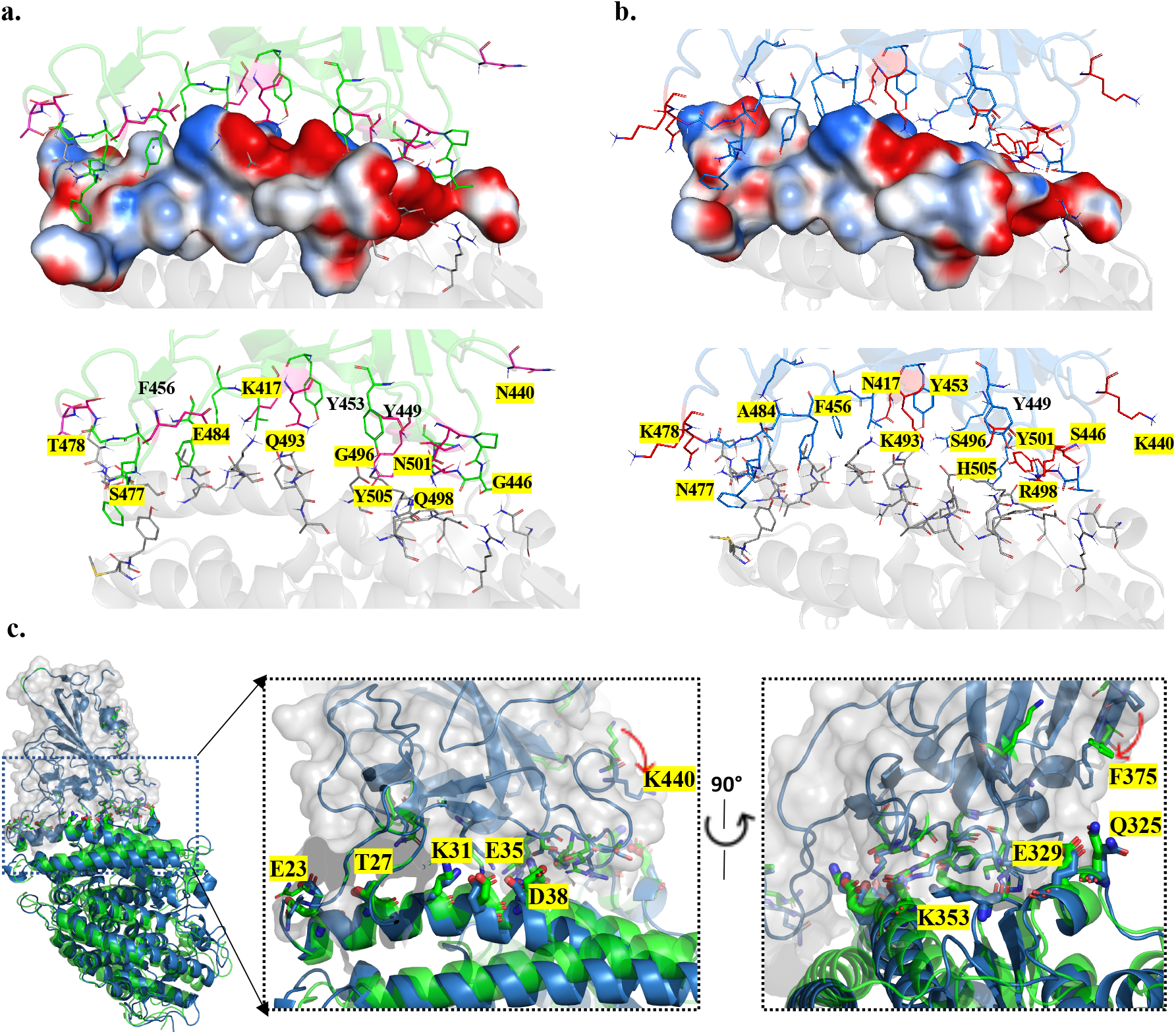
Detailed structures at the ACE2-RBD binding interface. (**a**) The interface of the wild type ACE2-RBD complex, the amino acids at mutation sites are shown with stick representations. The upper panels show the side chain positions of RBD on the surface of ACE2, where the surface is colored according to electrostatic potentials (−5 k_b_T/e to +5 k_b_T/e, for colors from red to blue); lower panel shows the side chains of both ACE2 and RBD. (**b**) The interactions between ACE2-RBDO of the Omicron variant. The figure labeling and coloring scheme are the same as in (a). The amino acids at mutation sites are highlighted with yellow color. (**c**) The conformation and the positions of ACE2 residues that are in close contact with RBD. The predicted complex model is shown in green color, and the representative model is in blue color. The RBD domain is enclosed by the solvent accessible surface colored in gray. The right panels show enlarged views of the interface in two orientations. The key residue side chains are shown in thicker sticks. Red arrows indicate the major movements of RBD residue side chains.

As a result of these mutations, wild type RBD-ACE2 interactions (Figure 5a) such as salt bridge E484-K31 are lost, K417-D30 are weakened in the Omicron variant due to shortened side chains, while hydrogen bonds Q493-E35, Q498-K353, Y505-E37 are enhanced by the Omicron substitutions, repositioning and forming new interactions, such as the favorable interactions K493-D38, R498-Y41, R498-Q42 H505-K353 and N377-Q24. Mutations also introduce additional π-π stacking interaction Y501-Y41. The key interactions observed in wild type ACE2-RBD are maintained in the Omicron variant (Figure 5b). These preserved interaction includes the following pairs: Y449-D38, Y453-H34 A475-S19, N487-Y83, T500-N330 and T500-D355.

Although the structures are highly similar in terms of backbone traces, there are notable conformational differences between the initial structure of the complex and the representative structure near the ACE2-RBD interface (Figure 5c). The N-terminal helices exhibit slightly kinked conformations, suggesting a larger separation from Omicron RBD by appearance. Nonetheless, careful analysis shows that the major binding interactions are well maintained through simulations, manifested as the highly consistent positions of key residues of ACE2 (highlighted in Figure 5c). The changes of RBD residue side chain positions suggest that MD simulations are useful in refining the quality of predicted complex structures. The side chains of F375 and K400 both point towards the ACE2 receptor in the representative structure, providing auxiliary supports to binding interactions (Figure 5c).

## Discussions and Conclusion

The large number of mutations observed in the spike protein of SARS-CoV-2 raised serious concerns about the new variant Omicron. Using computational modeling and simulations, we carried out quantitative analysis on the stability of ACE2-RBD complex for the Omicron variant, and compared to that of the wild type system. The interactions were assessed using several quantities, including hydrogen bonds, van der Waals contacts, buried surface areas, and the binding free energies. The dynamics simulation results and the quantitative comparison show that the binding interactions between ACE2 and RBD are slightly stronger for the Omicron variant than for the wild type. This information provides molecular basis for enhanced infectivity of the Omicron variant.

Most of effective neutralization antibodies are found to bind to RBD epitopes, many of them compete with ACE2 interactions, previous study has found that many of the neutralization antibodies are still effective to a large extend against the SARS CoV2 variants before Omicron variant [20,21]. However, the latest results have shown that 85% of previously characterized neutralization antibodies lost their efficacy against the new variant Omicron [22]. Therefore, the analyses of RBD-ACE2 interaction are not only important for the understanding of the outcome of the new virus variant, but also crucial for predicting and design for therapeutic antibody efficacy, particularly for further development of new generations of therapeutic antibodies that can overcome immune escaping mutants.

## Methods

### Molecular Dynamics Simulation and Analysis

The mutation information of Omicron is retrieved from the US CDC website [9]. We included 15 mutations occurred in the RDB (see Figure 1). The mutations were implemented based on the wild type ACE2-RBD complex structure using the Charmm-GUI webserver [23]. The protonation state was determined under PH 7.0 solvent environment.

The wild type ACE2-RBD and its Omicron variant were prepared using the CHARMM36 force fields, following the procedure of the CHARMM-GUI webserver. Each system was solvated in 150 mM sodium chloride solvent with TIP3P water models. Steepest descent algorithm was applied to minimize the system energy, then each system was equilibrated to 310.15 K (37 °C) within 125 ps. The temperature was maintained by Nose-Hoover scheme with 1.0 ps coupling constant in the NVT ensemble (constant volume and temperature). During the equilibration stage, harmonic restraint forces were applied to the molecules (400 kJ mol^-1^ nm^-2^ on backbone and 40 kJ mol^-1^ nm^-2^ on the side chain atoms) [24,25]. Subsequently, the harmonic restraints were removed and the NPT ensembles (constant pressure and temperature) were simulated at one atmosphere pressure (10^5^ Pa) and 310.15 K. The pressure was maintained by isotropic Parrinello-Rahman barostat [26], with a compressibility of 4.5 × 10^-5^ bar^-1^ and a coupling time constant of 5.0 ps. The wild type and Omicron variant ACE2-RBD systems were both simulated for 2 x 500 ns using the GROMACS 5.1.2 package [27]. In all simulations, a time step of 2.0 fs was used and the PME (particle mesh Ewald) [28] was applied for electrostatic interactions beyond 12.0 Å. The van der Waals interaction cutoff was set to 12.0 Å. Hydrogen atoms were constrained using the LINCS algorithm [29].

Analyses were carried out with tools in GROMACS (rmsd, rmsf, mindist, sasa) to examine the system stability. The buried surface area is computed as

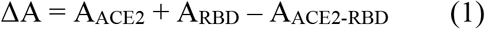

Where A_ACE2_, A_RBD_, and A_ACE2-RBD_ are the solvent accessible surface area computed using gmx sasa function. The mindist command was used to compute the residue distances, the residue pairs with distance below 4.0 Å were considered as contacting residues. VMD was used to analyze hydrogen bonding interactions [30], with the following criteria: D-A distance cutoff=3.9 Å and D-H-A angle cutoff=20 degrees, where D,A,H are Donor atom, Acceptor atom, and the Hydrogen atom linked to the Donor atom. Pymol was used for molecular binding interface, water distributions, visualization, and rending model images [30]. The adaptive Poisson-Boltzmann equation solver (APBS) was used to compute the electrostatic potentials [31].

The binding energy was calculated using Prime 3.0 MM-GBSA module of the Schrodinger 24 package [32–34]. In each ACE2-RBD complex, the ACE2 was treated as the receptor and RBD was considered as the ligand. Prime MM-GBSA uses OPLS-AA force field and VSGB 2.0 implicit solvation model to estimate the binding energy of the receptor-ligand complex. The binding energy is calculated as:

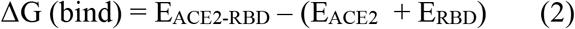

## Acknowledgement

This work was supported by the National Key Research and Development Projects of the Ministry of Science and Technology of China (2021YFC2301300) to X.-D.S, and the National Natural Science Foundation of China (grant numbers: U1930402, 31971136) to H.L. The computational work is supported by a Tianhe-2JK computing time award at Beijing Computational Science Research Center (CSRC).

## Competing interests

The authors declare no competing interests.

## Notes

### Competing Interest Statement

The authors have declared no competing interest.

### Summary of Updates

The figure 5 is updated. There is a mistake in the original figure 4, where the percentage values were swapped for the wild type and Omicron systems. There is improvement in grammar and wording.

## Reference

[1] I. Ishigami, N.A. Zatsepin, M. Hikita, C.E. Conrad, G. Nelson, J.D. Coe, S. Basu, T.D. Grant, M.H. Seaberg, R.G. Sierra, M.S. Hunter, P. Fromme, R. Fromme, S.R. Yeh, D.L. Rousseau, Crystal structure of CO-bound cytochrome c oxidase determined by serial femtosecond X-ray crystallography at room temperature, Proc. Natl. Acad. Sci. U. S. A. 114 (2017) 8011–8016. https://doi.org/10.1073/pnas.1705628114.

[2] Johns Hopkins Coronavirus Resource Center, (n.d.). https://coronavirus.jhu.edu/vaccines/vaccines-faq (accessed December 13, 2021).

[3] E. Mathieu, H. Ritchie, E. Ortiz-Ospina, M. Roser, J. Hasell, C. Appel, C. Giattino, L. Rodés-Guirao, A global database of COVID-19 vaccinations, Nat. Hum. Behav. 5 (2021) 947–953. https://doi.org/10.1038/s41562-021-01122-8.

[4] J.S. Tregoning, K.E. Flight, S.L. Higham, Z. Wang, B.F. Pierce, Progress of the COVID-19 vaccine effort: viruses, vaccines and variants versus efficacy, effectiveness and escape, Nat. Rev. Immunol. 2021 2110. 21 (2021) 626–636. https://doi.org/10.1038/s41577-021-00592-1.

[5] P.R. Krause, T.R. Fleming, R. Peto, I.M. Longini, J.P. Figueroa, J.A.C. Sterne, A. Cravioto, H. Rees, J.P.T. Higgins, I. Boutron, H. Pan, M.F. Gruber, N. Arora, F. Kazi, R. Gaspar, S. Swaminathan, M.J. Ryan, A.M. Henao-Restrepo, Considerations in boosting COVID-19 vaccine immune responses, Lancet. 398 (2021) 1377–1380. https://doi.org/10.1016/S0140-6736(21)02046-8/ATTACHMENT/99B4689E-FA37-476A-B30E-8D1E3B4E33BCZMMC1.PDF.

[6] A.P. Vashi, O.C. Coiado, The future of COVID-19: A vaccine review, J. Infect. Public Health. 14 (2021) 1461–1465. https://doi.org/10.1016/J.JIPH.2021.08.011.

[7] L. Wang, T. Zhou, Y. Zhang, E.S. Yang, C.A. Schramm, W. Shi, A. Pegu, O.K. Oloniniyi, A.R. Henry, S. Darko, S.R. Narpala, C. Hatcher, D.R. Martinez, Y. Tsybovsky, E. Phung, O.M. Abiona, A. Antia, E.M. Cale, L.A. Chang, M. Choe, K.S. Corbett, R.L. Davis, A.T. DiPiazza, I.J. Gordon, S.H. Hait, T. Hermanus, P. Kgagudi, F. Laboune, K. Leung, T. Liu, R.D. Mason, A.F. Nazzari, L. Novik, S. O’Connell, S. O’Dell, A.S. Olia, S.D. Schmidt, T. Stephens, C.D. Stringham, C.A. Talana, I.T. Teng, D.A. Wagner, A.T. Widge, B. Zhang, M. Roederer, J.E. Ledgerwood, T.J. Ruckwardt, M.R. Gaudinski, P.L. Moore, N.A. Doria-Rose, R.S. Baric, B.S. Graham, A.B. McDermott, D.C. Douek, P.D. Kwong, J.R. Mascola, N.J. Sullivan, J. Misasi, Ultrapotent antibodies against diverse and highly transmissible SARS-CoV-2 variants, Science. 373 (2021). https://doi.org/10.1126/science.abh1766.

[8] Classification of Omicron (B.1.1.529): SARS-CoV-2 Variant of Concern, (n.d.). https://www.who.int/news/item/26-11-2021-classification-of-omicron-(b.1.1.529)-sars-cov-2-variant-of-concern (accessed December 9, 2021).

[9] Science Brief: Omicron (B.1.1.529) Variant | CDC, (n.d.). https://www.cdc.gov/coronavirus/2019-ncov/science/science-briefs/scientific-brief-omicron-variant.html (accessed December 9, 2021).

[10] J. Shang, G. Ye, K. Shi, Y. Wan, C. Luo, H. Aihara, Q. Geng, A. Auerbach, F. Li, Structural basis of receptor recognition by SARS-CoV-2, Nature. 581 (2020) 221–224. https://doi.org/10.1038/s41586-020-2179-y.

[11] R. Yan, Y. Zhang, Y. Li, L. Xia, Y. Guo, Q. Zhou, Structural basis for the recognition of SARS-CoV-2 by full-length human ACE2, Science. 367 (2020) 1444–1448. https://doi.org/10.1126/science.abb2762.

[12] Y. Wang, M. Liu, J. Gao, Enhanced receptor binding of SARS-CoV-2 through networks of hydrogen-bonding and hydrophobic interactions, Proc. Natl. Acad. Sci. 117 (2020) 13967–13974. https://doi.org/10.1073/pnas.2008209117.

[13] C.S. Lupala, X. Li, J. Lei, H. Chen, J. Qi, H. Liu, X.-D. Su, Computational simulations reveal the binding dynamics between human ACE2 and the receptor binding domain of SARS-CoV-2 spike protein, Quant. Biol. 9 (2021) 61. https://doi.org/10.15302/J-QB-020-0231.

[14] X. Xu, P. Chen, J. Wang, J. Feng, H. Zhou, X. Li, W. Zhong, P. Hao, Evolution of the novel coronavirus from the ongoing Wuhan outbreak and modeling of its spike protein for risk of human transmission, Sci. China Life Sci. 63 (2020) 457–460. https://doi.org/10.1007/s11427-020-1637-5.

[15] J. Damas, G.M. Hughes, K.C. Keough, C.A. Painter, N.S. Persky, M. Corbo, M. Hiller, K.P. Koepfli, A.R. Pfenning, H. Zhao, D.P. Genereux, R. Swofford, K.S. Pollard, O.A. Ryder, M.T. Nweeia, K. Lindblad-Toh, E.C. Teeling, E.K. Karlsson, H.A. Lewin, Broad host range of SARS-CoV-2 predicted by comparative and structural analysis of ACE2 in vertebrates, Proc. Natl. Acad. Sci. U. S. A. 117 (2020) 22311–22322. https://doi.org/10.1073/pnas.2010146117.

[16] C.S. Lupala, V. Kumar, X. Su, C. Wu, H. Liu, Computational insights into differential interaction of mammalian angiotensin-converting enzyme 2 with the SARS-CoV-2 spike receptor binding domain, Comput. Biol. Med. (2021) 105017. https://doi.org/10.1016/j.compbiomed.2021.105017.

[17] C. Yi, X. Sun, J. Ye, L. Ding, M. Liu, Z. Yang, X. Lu, Y. Zhang, L. Ma, W. Gu, A. Qu, J. Xu, Z. Shi, Z. Ling, B. Sun, Key residues of the receptor binding motif in the spike protein of SARS-CoV-2 that interact with ACE2 and neutralizing antibodies, Cell. Mol. Immunol. 17 (2020) 621–630. https://doi.org/10.1038/s41423-020-0458-z.

[18] Q. Wang, Y. Zhang, L. Wu, S. Niu, C. Song, Z. Zhang, G. Lu, C. Qiao, Y. Hu, K.Y. Yuen, Q. Wang, H. Zhou, J. Yan, J. Qi, Structural and Functional Basis of SARS-CoV-2 Entry by Using Human ACE2, Cell. 181 (2020) 894–904.e9. https://doi.org/10.1016/j.cell.2020.03.045.

[19] J. Lan, J. Ge, J. Yu, S. Shan, H. Zhou, S. Fan, Q. Zhang, X. Shi, Q. Wang, L. Zhang, X. Wang, Structure of the SARS-CoV-2 spike receptor-binding domain bound to the ACE2 receptor., Nature. (2020). https://doi.org/10.1038/s41586-020-2180-5.

[20] S. Du, P. Liu, Z. Zhang, T. Xiao, A. Yasimayi, W. Huang, Y. Wang, Y. Cao, X.S. Xie, J. Xiao, Structures of SARS-CoV-2 B.1.351 neutralizing antibodies provide insights into cocktail design against concerning variants, Cell Res. 31 (2021) 1130–1133. https://doi.org/10.1038/s41422-021-00555-0.

[21] H. Xu, B. Wang, T.N. Zhao, Z.T. Liang, T.B. Peng, X.H. Song, J.J. Wu, Y.C. Wang, X.D. Su, Structure-based analyses of neutralization antibodies interacting with naturally occurring SARS-CoV-2 RBD variants, Cell Res. 31 (2021) 1126–1129. https://doi.org/10.1038/s41422-021-00554-1.

[22] Y. Cao, J. Wang, F. Jian, T. Xiao, W. Song, A. Yisimayi, W. Huang, Q. Li, P. Wang, R. An, J. Wang, Y. Wang, X. Niu, S. Yang, H. Liang, H. Sun, T. Li, Y. Yu, Q. Cui, S. Liu, X. Yang, S. Du, Z. Zhang, X. Hao, F. Shao, R. Jin, X. Wang, J. Xiao, Y. Wang, X.S. Xie, B.1.1.529 escapes the majority of SARS-CoV-2 neutralizing antibodies of diverse epitopes, BioRxiv. (2021) 2021.12.07.470392. https://doi.org/10.1101/2021.12.07.470392.

[23] J. Lee, X. Cheng, J.M. Swails, M.S. Yeom, P.K. Eastman, J.A. Lemkul, S. Wei, J. Buckner, J.C. Jeong, Y. Qi, S. Jo, V.S. Pande, D.A. Case, C.L. Brooks, A.D. MacKerell, J.B. Klauda, W. Im, CHARMM-GUI Input Generator for NAMD, GROMACS, AMBER, OpenMM, and CHARMM/OpenMM Simulations Using the CHARMM36 Additive Force Field, J. Chem. Theory Comput. 12 (2016) 405–413. https://doi.org/10.1021/acs.jctc.5b00935.

[24] S. Nosé, A unified formulation of the constant temperature molecular dynamics methods, J. Chem. Phys. 81 (1984) 511–519. https://doi.org/10.1063/1.447334.

[25] W.G. Hoover, Canonical dynamics: Equilibrium phase-space distributions, Phys. Rev. A. 31 (1985) 1695–1697. https://doi.org/10.1103/PhysRevA.31.1695.

[26] M. Parrinello, A. Rahman, Polymorphic transitions in single crystals: A new molecular dynamics method, J. Appl. Phys. 52 (1981) 7182–7190. https://doi.org/10.1063/1.328693.

[27] M.J. Abraham, T. Murtola, R. Schulz, S. Páll, J.C. Smith, B. Hess, E. Lindah, Gromacs: High performance molecular simulations through multi-level parallelism from laptops to supercomputers, SoftwareX. 1–2 (2015) 19–25. https://doi.org/10.1016/j.softx.2015.06.001.

[28] T. Darden, D. York, L. Pedersen, Particle mesh Ewald: An N·log(N) method for Ewald sums in large systems, J. Chem. Phys. 98 (1993) 10089–10092. https://doi.org/10.1063/1.464397.

[29] B. Hess, H. Bekker, H.J.C. Berendsen, J.G.E.M. Fraaije, LINCS: A Linear Constraint Solver for molecular simulations, J. Comput. Chem. 18 (1997) 1463–1472. https://doi.org/10.1002/(SICI)1096-987X(199709)18:12<1463::AID-JCC4>3.0.CO;2-H.

[30] W. Humphrey, A. Dalke, K. Schulten, VMD: Visual molecular dynamics, J. Mol. Graph. 14 (1996) 33–38. https://doi.org/10.1016/0263-7855(96)00018-5.

[31] E. Jurrus, D. Engel, K. Star, K. Monson, J. Brandi, L.E. Felberg, D.H. Brookes, L. Wilson, J. Chen, K. Liles, M. Chun, P. Li, D.W. Gohara, T. Dolinsky, R. Konecny, D.R. Koes, J.E. Nielsen, T. Head-Gordon, W. Geng, R. Krasny, G.W. Wei, M.J. Holst, J.A. McCammon, N.A. Baker, Improvements to the APBS biomolecular solvation software suite, Protein Sci. 27 (2018) 112–128. https://doi.org/10.1002/pro.3280.

[32] Schrödinger LLC, Schrödinger Release 2021-4: Prime, (2021).

[33] M.P. Jacobson, D.L. Pincus, C.S. Rapp, T.J.F. Day, B. Honig, D.E. Shaw, R.A. Friesner, A hierarchical approach to all-atom protein loop prediction, Proteins Struct. Funct. Bioinforma. 55 (2004) 351–367. https://doi.org/10.1002/PROT.10613.

[34] M.P. Jacobson, R.A. Friesner, Z. Xiang, B. Honig, On the Role of the Crystal Environment in Determining Protein Side-chain Conformations, J. Mol. Biol. 320 (2002) 597–608. https://doi.org/10.1016/S0022-2836(02)00470-9.

